# Inbred rat heredity and sex affect oral oxycodone self-administration and augmented intake in long sessions: correlations with anxiety and novelty-seeking

**DOI:** 10.1101/2023.11.26.568753

**Authors:** Burt M Sharp, Shuangying Leng, Jun Huang, Caroline Jones, Hao Chen

## Abstract

Oxycodone abuse begins with prescription oral oxycodone, yet vulnerability factors determining abuse are largely undefined. We evaluated genetic vulnerability in a rat model of oral oxycodone self-administration (SA): increasing oxycodone concentration/session (0.025-0.1mg/ml; 1,4,16-h) followed by extinction and reinstatement. Active licks and oxycodone intake were greater in females than males during 4-h and 16-h sessions (p< 0.001). Each sex increased intake during 16-h vs 4-h sessions (p<2e-16), but a subset of strains dramatically augmented intake at 16-h (p=0.0005). Heritability (*h*^2^) of active licks/4-h at increasing oxycodone dose ranged from 0.30-0.53. Under a progressive ratio schedule, breakpoints were strain-dependent (p<2e-16). Cued reinstatement was greater in females (p<0.001). Naive rats were assessed by elevated plus maze (EPM), open field (OF) and novel object interaction (NOI). We correlated these behaviors with 28 parameters of oxycodone SA. Anxiety-defining EPM traits were most associated with SA in both sexes, whereas more OF and NOI traits were SA-associated in males. Sex and heredity are major determinants of motivation to take and seek oxycodone; intake augments dramatically during extended access in specific strains; and pleiotropic genes affect anxiety and multiple SA parameters.

## Introduction

Opioids such as morphine and *oxycodone* have long been prescribed for effective analgesia. However, the prevalence of prescriptions for opioids such as oxycodone, has led to widespread abuse and deaths. Approximately 12 million people misused opioids in 2016 ^1^. By 2017, deaths from opioid overdoses rose to approximately 12 million, and in 2022, 80,000 opioid-involved overdose deaths were reported ^2^.

Most commonly, individuals initiate their habitual intake of oxycodone with prescription oral oxycodone ^3,4^. Since pharmacokinetic parameters are important determinants of abuse potential, we designed a rat oral operant self-administration (SA) model. Rats were used to model the oral pattern of drug intake common in humans. Previous reports of oral oxycodone SA ^5–7^ required training procedures to initiate drug SA that alter the motivation to obtain drugs. The model we developed ^8^ utilizes operant licking for oxycodone and does not require water restriction or prior drug exposure.

The transition from deliberately controlled drug intake to compulsive drug seeking and taking, characteristic of addiction ^9^, is often accompanied by a sharp rise in drug use. The marked variation in amount of drug consumed and the extent of addictive behavior observed amongst individuals depends on multiple factors. In human twin studies, approximately half of the vulnerability to develop an opiate use disorder (OUD) is heritable ^10,11^. Complex, sex-dependent patterns also determine opiate use. Male rats self-administer more oxycodone than females at early stages, while females are more susceptible to opioid reward ^12^.

Based on the known impact of genetics and sex on the vulnerability to OUD, the purpose of these studies was to identify the genetically determined behavioral parameters of oral oxycodone self-administration. To accomplish this, we studied the patterns of drug intake in both sexes during 4-h and 16-h extended sessions; a total of 36 fully inbred rat strains were evaluated, 23 were common to both sexes. All inter-sex statistical comparisons were restricted to the 23 common strains. We also correlated oxycodone SA parameters to exploratory behaviors and locomotion in drug naive rats.

Some strains manifest greater than a 100-fold *increase in oxycodone intake* during our 65-day protocol. We also found significant sex and strain differences that affect multiple parameters of oxycodone SA, including the dramatic strain-specific augmentation of intake with increased drug availability during 16-h access sessions and the correlation of intake between 4 vs 16-h sessions in both sexes. We confirmed the significant heritability (*h^2^*) of multiple parameters of oxycodone SA including the motivation to take the drug, the amount consumed, the behavior emitted (i.e., active licks) to obtain drug, and the reinstatement of drug seeking behavior. These findings are similar to those reported in seven inbred rat strains ^8^ and in studies of human twins. Lastly, we found significant sex and strain-dependent correlations between specific behavioral tests (e.g., elevated plus maze, EPM) conducted in drug naive rats and oxycodone SA. Overall, this oral model of oxycodone SA captures multiple behavioral parameters involved in the strong abuse potential of oral oxycodone and demonstrates the existence of pleiotropic genes shared between these behavioral parameters of oxycodone SA and independent behaviors elicited in paradigms such as EPM.

## Results

### Oxycodone SA in Males and Females

Our operant conditioning protocol reinforces the licking of a drug delivery spout by delivering one drop (60 μl) of oxycodone under a fixed ratio 5 (FR5, with a 20s time out period) schedule. The ∼65 day procedure is summarized in Table 1 of Methods.

**Table 1.**
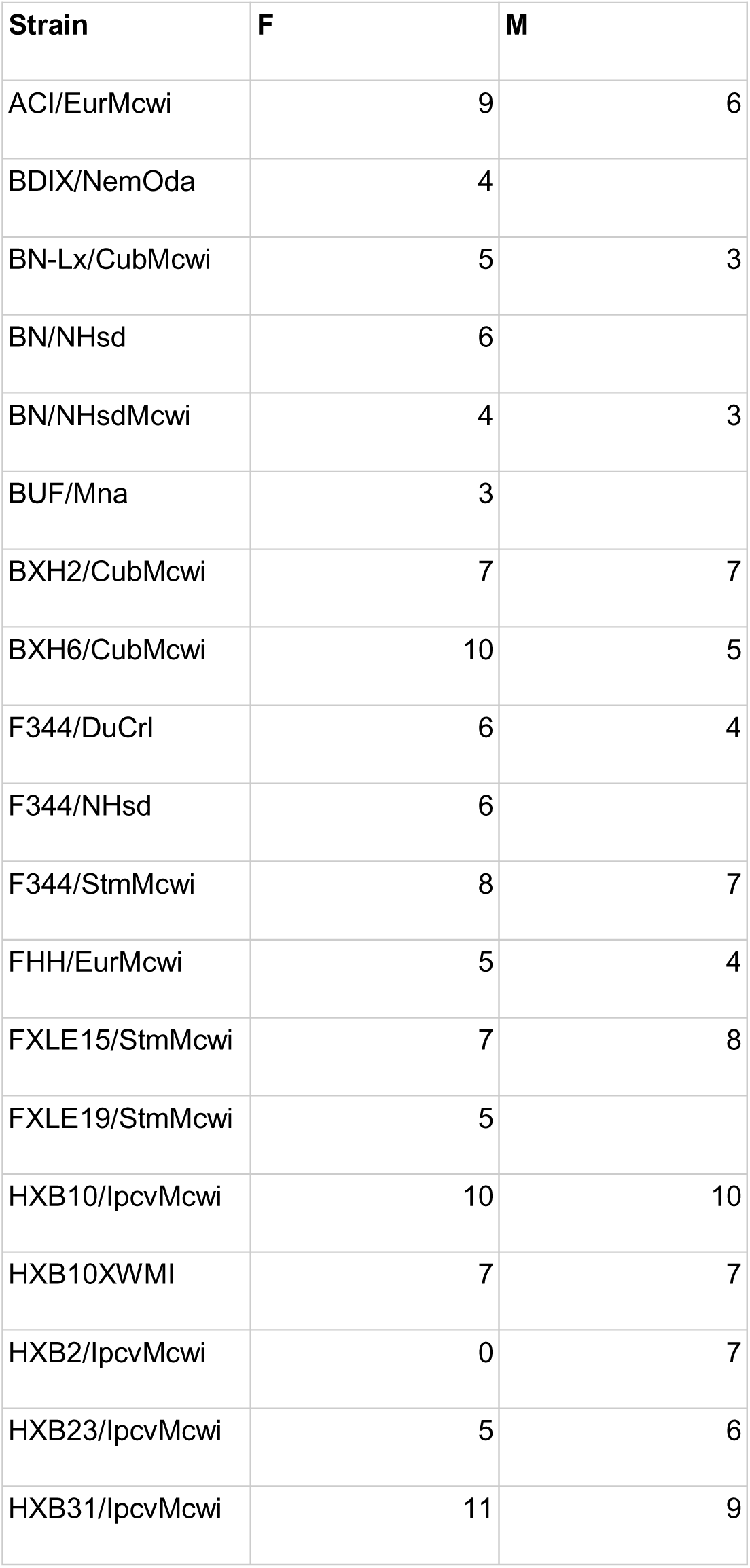
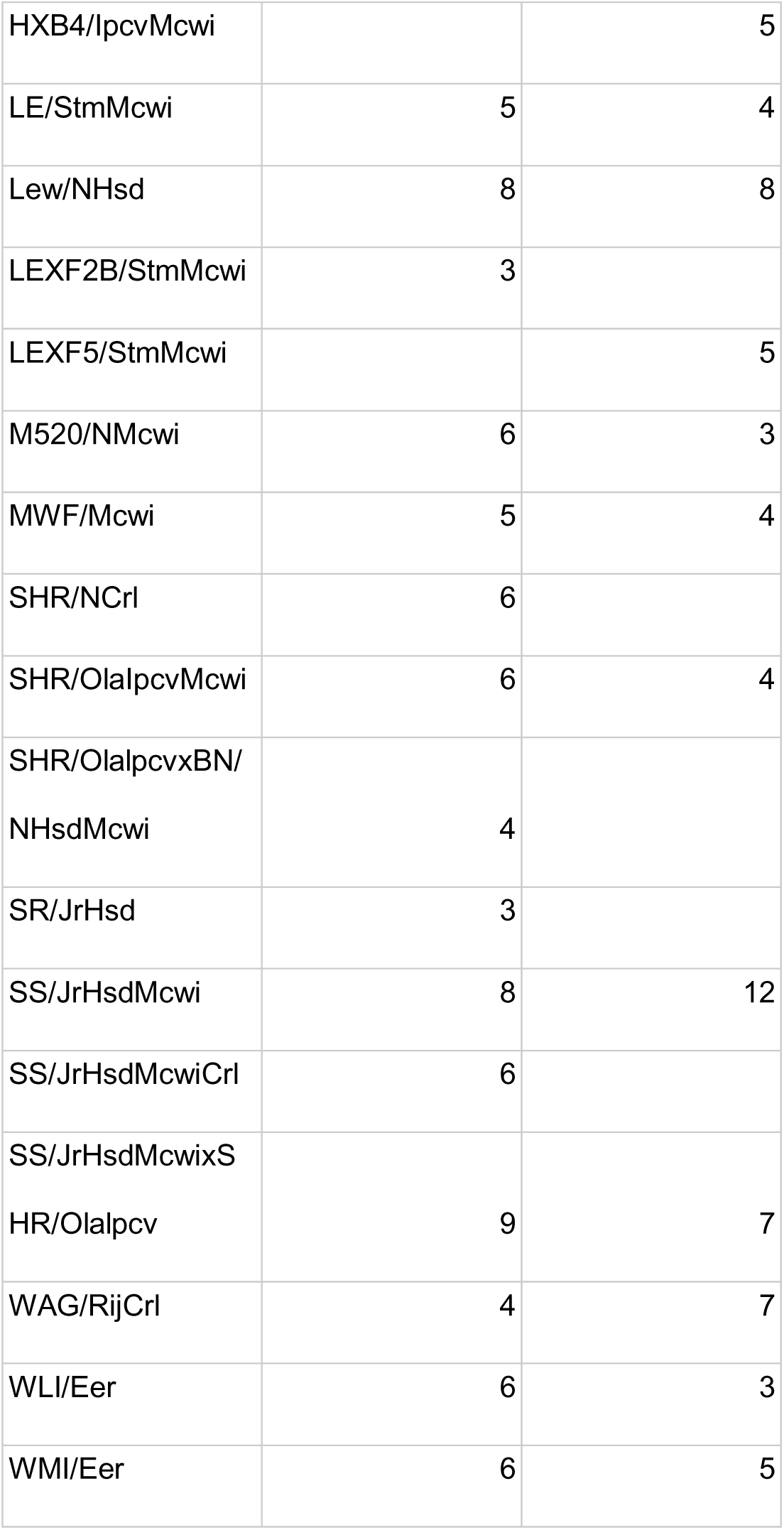
The number of rats per strain for each sex.

Of 36 strains included in the hybrid rat diversity panel (HRDP; ^13^), oral operant oxycodone SA was studied in females and males from 33 and 26 strains, respectively (Figure 2A,B). The number of active (mean±sem) and inactive licks emitted by rats across all strains are shown for operant sessions that increased in duration (1-h/4-h/16-h) and dose (0.025-0.1 mg/kg) followed by 4 extinction sessions on alternating days and reinstatement of extinguished drug seeking behavior. A progressive ratio schedule of oxycodone 0.1 mg/kg was conducted on the day prior to the first 16-h session. In both sexes (Figure 2), a strong preference for the active vs. inactive spout was observed across the HRDP at all doses and session durations. In the 23 strains common to both sexes, active licks were greater in females than males during both 4-h and 16-h sessions (panel C: 4-h, F_1,275_=9.14, p=0.003; 16-h, F_1,233_=4.97, p=0.03). Inactive licks were not different between sexes (panel D: 4-h, F_1,275_=2.7, p>0.05; 16-h, F_1,233_=0.76, p>0.05).

**Figure 1.**
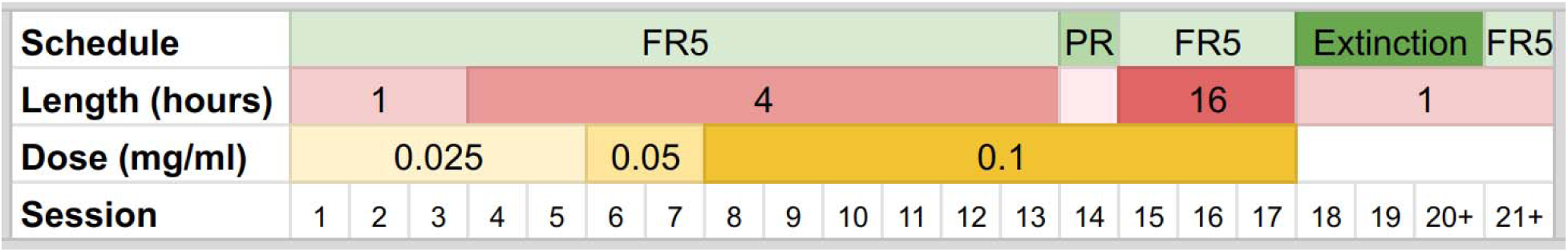
Protocol and schedule of operant oxycodone self-administration. The first three sessions were conducted daily. Subsequent sessions were conducted on alternate days. The final scheduled FR5 session was the reinstatement of extinguished oxycodone seeking.

**Figure 2.**
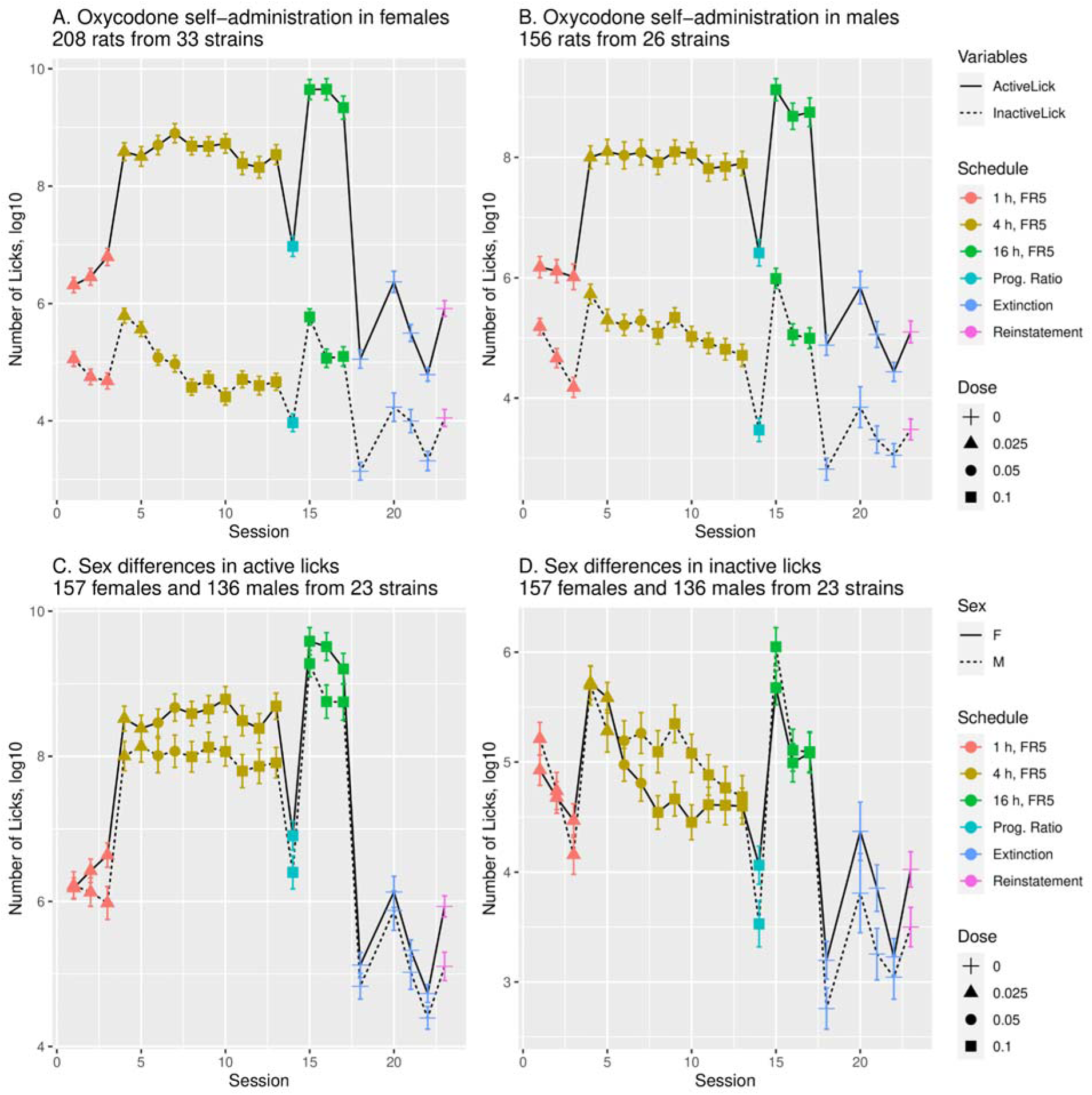
Number of licks during oxycodone SA. Rats showed a strong preference for the active over inactive spout in females (A) and males (B) throughout the HRDP. (C) The number of licks on the active spout was greater in females (n=157) than in males (n=136) during both 4-h (p=0.003) and 16-h sessions (p=0.03) in the 23 strains where both sexes were studied. (D) The number of licks on the inactive spout was not different between sexes (p>0.05) during either the 4-h or 16-h sessions.

There was a strong sex difference in oxycodone intake across the 23 strains common to both sexes (Figure 3). In females, across all SA sessions, mean rewards/session (panel A: F_1,275_=125.92, p<2e-16) and intake/session (B: F_1,275_=105.23, p<2e-16) were greater than in males. Similarly, there were sex differences in oxycodone intake across different SA stages (panel C), including 1-h sessions (F_1,275_=66.8, p=1.1e-14), 4-h sessions (F_1,275_=119.9, p<2e-16), and 16-h sessions (F_1,233_=79.7, p<2e16). In both sexes given access to the same dose of oxycodone (0.1 mg/ml) during 4-h and 16-h sessions, the greatest intake occurred during the long 16-h sessions: intake increased in both males (F_1,753_=83.6, p=2e-16) and females (F_1,901_=162.9, p=2e-16).

**Figure 3.**
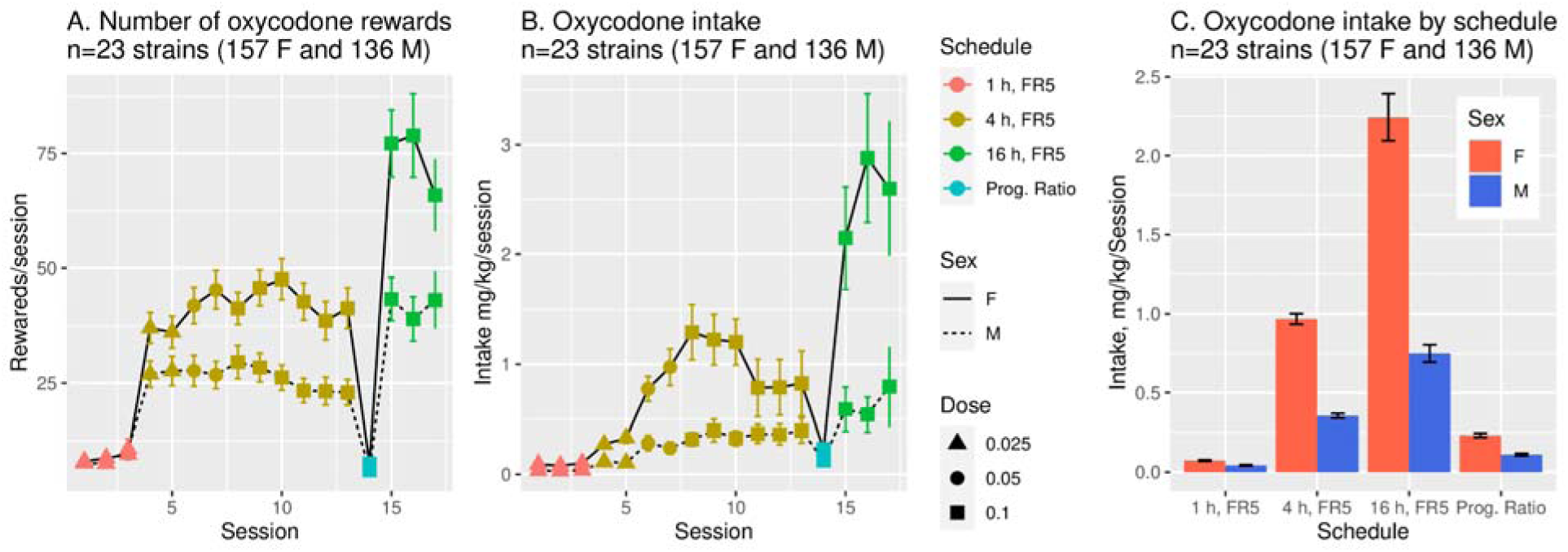
Sex differences in oxycodone rewards and intake. The mean oxycodone reward (A) and intake (B) per session (FR5) were significantly greater in females than males in the 23 strains where both sexes were studied (p<0.001 for both). Intake was also significantly greater in females than in males across different stages (C) of SA (p<0.001 for all). In both sexes, the greatest intake occurred during 16-h sessions.

Across the 36 HRDP strains, oxycodone intake was strain-dependent in both sexes (Figure 4). Mean intake during the first three 1-h sessions, when SA of oxycodone (0.025 mg/ml) was acquired (panels A. B), varied greatly by strain (male: F_25,432_=9.07, p=9.5e-27. female: F_32,570_=7.45, p=5.3e-27). Similarly, mean intake during the final three 4-h sessions, when stable oxycodone (0.1 mg/ml) intake was attained (panels C, D), varied greatly by strain in both sexes (F_25,506_=25.74, p=3.8e-74 in males and F_32,620_=9.59, p=1.30e-36).

**Figure 4.**
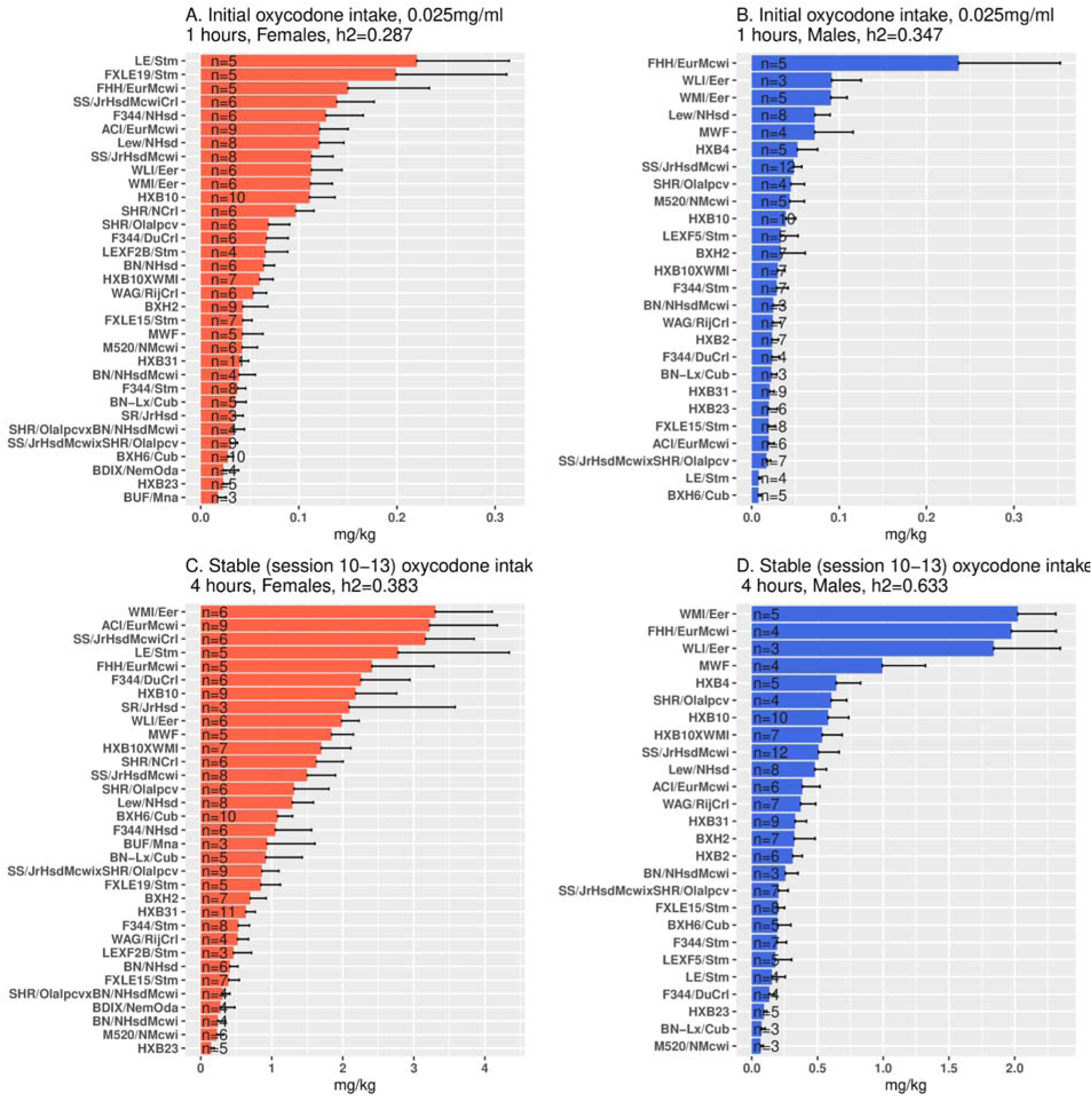
Initial 1-h and stable 4-h oxycodone intake within each sex across the 36 HRDP strains. Mean intake during the first three sessions (top 2 panels) showed large strain differences in each sex. Similarly, mean intake during the last three 4-h sessions (stable intake; lower 2 panels) varied across the HRDP in each sex. (note: the difference in X axis scales)

### Stable Oxycodone Intake during 4-h Limited Access vs.16-h Extended Access Sessions

We compared and correlated stable oxycodone intake (0.1 mg/ml) during limited 4-h vs. extended 16-h sessions within each sex across the 36 HRDP strains. Oxycodone intake in 16-h sessions exceeded 4-h sessions in many strains (Figure 5A, 5B). This change was sex-specific in that specific strains showed increased 16-h consumption in only one sex of the 23 common strains. In addition, a greater number of strains throughout HRDP showed increased 16-h oxycodone intake in males compared to females. Figures 5C and 5D show the correlations within females (r=0.832, p<0.0001) and within males (r=0.835, p<0.0001) respectively, for oxycodone intake in 4-h vs. 16-h sessions across all strains. In summary, in both sexes across the HRDP, oxycodone intake during 4-h sessions was predictive of intake during 16-h sessions. Within sex, the fold-increase in oxycodone intake during our 65-day SA protocol, from the initial 1-h sessions at 0.025 mg/ml to the final 16-h sessions at 0.1 mg/ml, varied greatly by strain. In males, the increase ranged from 4.67 (BN-Lx/CubMcwi) to 126.0-fold (LE/StmMcwi); in females, it was 7.2 (BN/NHsdMcwi) to 167.2-fold (SR/JrHsdMcwi).

**Figure 5.**
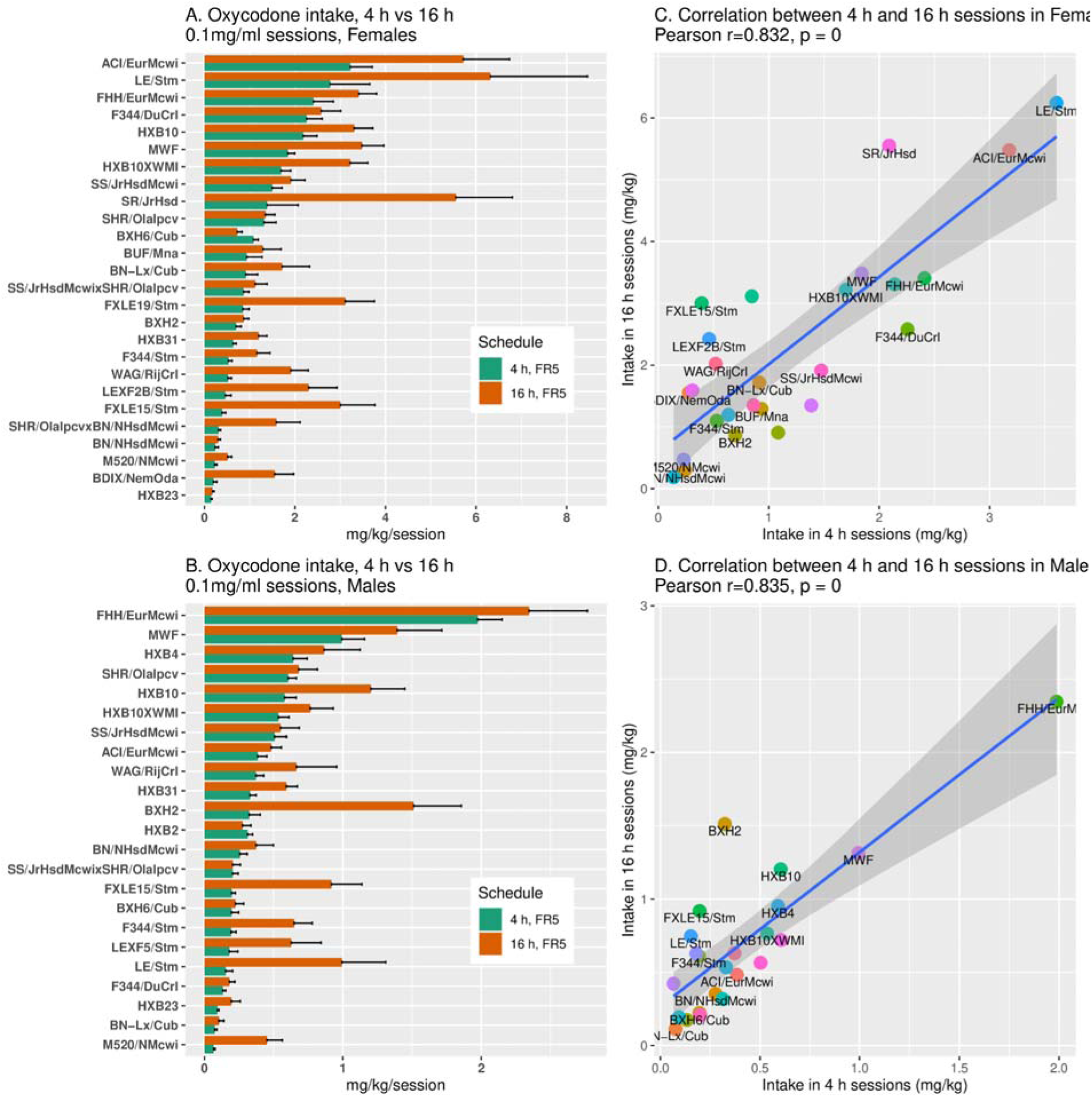
Oxycodone (0.1 mg/kg) intake in 4-h and 16-h sessions within each sex. A, B: Extending access from 4-h to 16-h increased total drug intake/session in many strains and both sexes. Within specific strains, increased 16-h intake was sex-specific. C, D: The correlation of oxycodone intake between 4 h and 16 h sessions was significant in both females (r=0.832, p< 0.0001) and males (r=0.835, p<0.0001).

### Augmented Oxycodone Intake During Long Access Sessions in a Subset of Strains

Across all strains, each sex increased intake during 16-h vs 4-h sessions (p<2e-16), but a subset of the 23 strains dramatically augmented (>3-fold) their intake at 16-h (Fig 6A, B; F_1,12_=15.03, p=0.002 for females and F_1,10_=18.22, p=0.002 for males). In contrast, oxycodone intake at 4-h was similar in augmenters vs non-augmenters within each sex (Fig 6A, B; female, F_1,12_=0.05, p=0.84; male, F_1,10_=0.08, p=0.79). Inter-sex comparisons were not conducted because only 3 strains were common to both sexes. Figure 6C-F is a high time resolution view of oxycodone intake by hour and sex during 4-h and 16-h sessions. In 4-h sessions (Figure 6C, D), maximum oxycodone intake occurred in both sexes of augmenters and non-augmenters during the first hour and declined thereafter toward baseline levels by 3-4-h. In 16-h sessions (Figure 6E, F), intake in non-augmenters was maximal at 1-h, declining gradually thereafter in both sexes. In contrast, early intake, from 1-6-h, was much greater in the augmenters of both sexes, rapidly decreasing from approximately 7-11-h, and gradually thereafter. By 13-16-h, intake within sex was similar in both augmenters and non-augmenters. Augmenters of both sexes have a prolonged interval of loading-up with oxycodone from approximately 1-6-h that is not observed in non-augmenters. Following this interval, oxycodone intake declines steadily, devoid of periodic increases.

**Figure 6.**
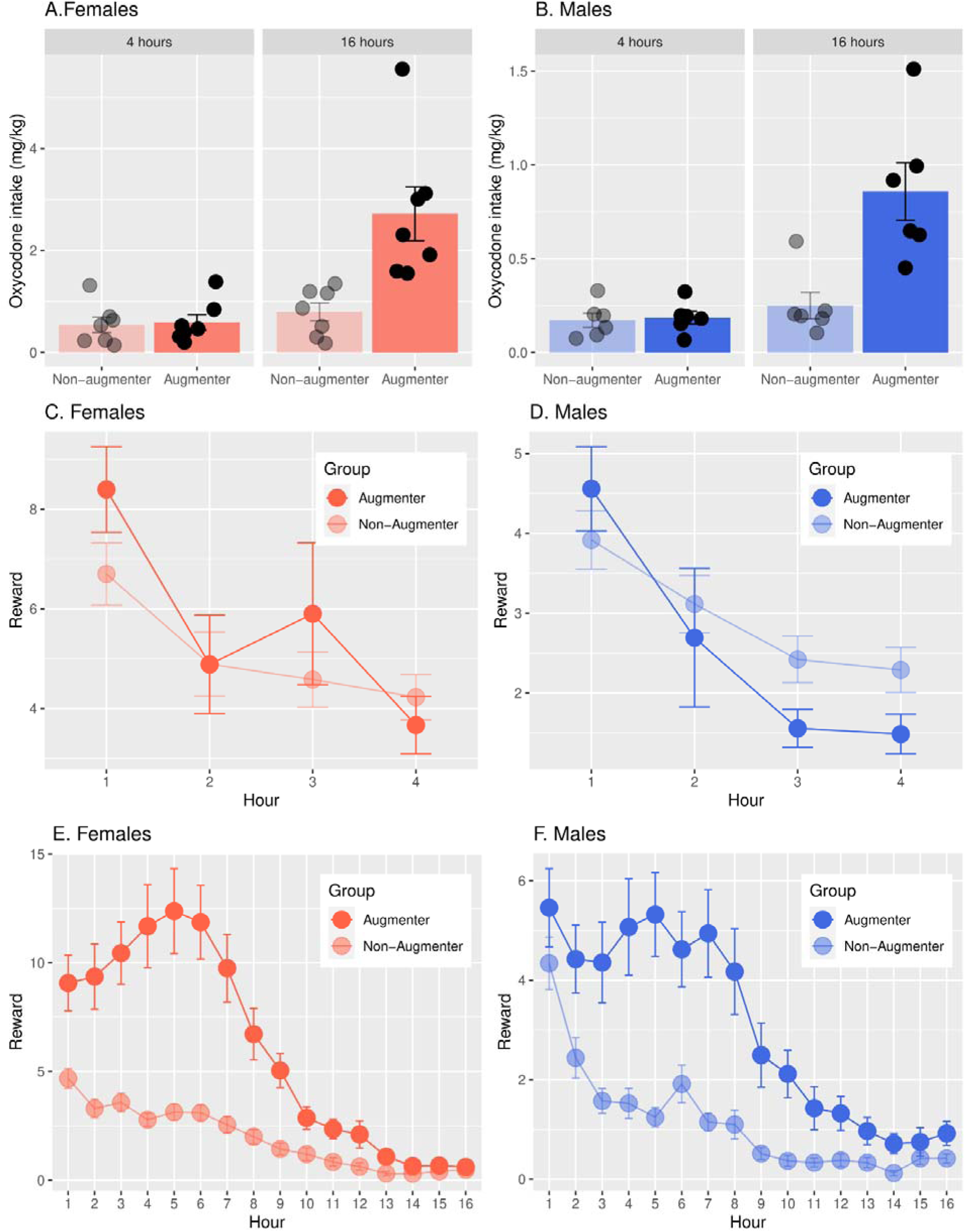
Augmented oxycodone intake in 16-h vs 4-h sessions. A subset of strains (7 female, 6 male) augmented their oxycodone intake during long access 16-h vs 4-h sessions by at least 3-fold (Females: F_1,12_=15.03, p=0.002; Males: F_1,10_=18.22, p=0.002, panels B). Inter-sex comparisons were not conducted because only 3 strains were common to both sexes. At 4-h, non-augmenters and augmenters had similar oxycodone intake within each sex (Females; F_1,12_=0.05, p=0.84, panels A; Males: F_1,10_=0.08, p=0.79, Panel B). In the 4-h sessions (panels C, D), maximum oxycodone intake occurred in both sexes of augmenters and non-augmenters during the first hour and declined thereafter toward baseline levels by 3-4 hours. In the 16-h sessions (panels E, F), intake in non-augmenters was maximal at 1 hour, declining gradually thereafter in both sexes. In contrast, early intake, from 1-6 hours, was much greater in the augmenters of both sexes, rapidly decreasing from approximately 7-11 hours, and gradually thereafter.

To determine whether genetic differences in oxycodone metabolism might contribute to the divergent patterns of 16-h oxycodone intake in augmenters vs non-augmenters, we identified 1,263 SNPs in different alleles associated with the 3 major metabolic enzymes that differed between these two groups. The allele frequencies of these SNPs were not different statistically, even at 30% false discovery rate. While these data strongly suggest that genetic variation is unlikely to affect the function of principal isoenzymes regulating oxycodone metabolism, they do not eliminate the possibility of variation in gene expression by other mechanisms (e.g., transcriptional or post-translational).

### Progressive Ratio Schedule and Reinstatement of Extinguished Oxycodone Seeking

In Figure 7, the breakpoints achieved were strain-dependent in each sex (A. females, F_32,148_=5.2, p=1.8e-12; B. males, F_25,103_=2.6, p=0.0003). In the subset of 23 strains where PR was measured in both sexes, the effect of sex was also significant (F_22,227_=6.4, p=2.5e-14). Therefore, in both sexes, the motivation to obtain oral oxycodone is strain- and sex-dependent.

**Figure 7.**
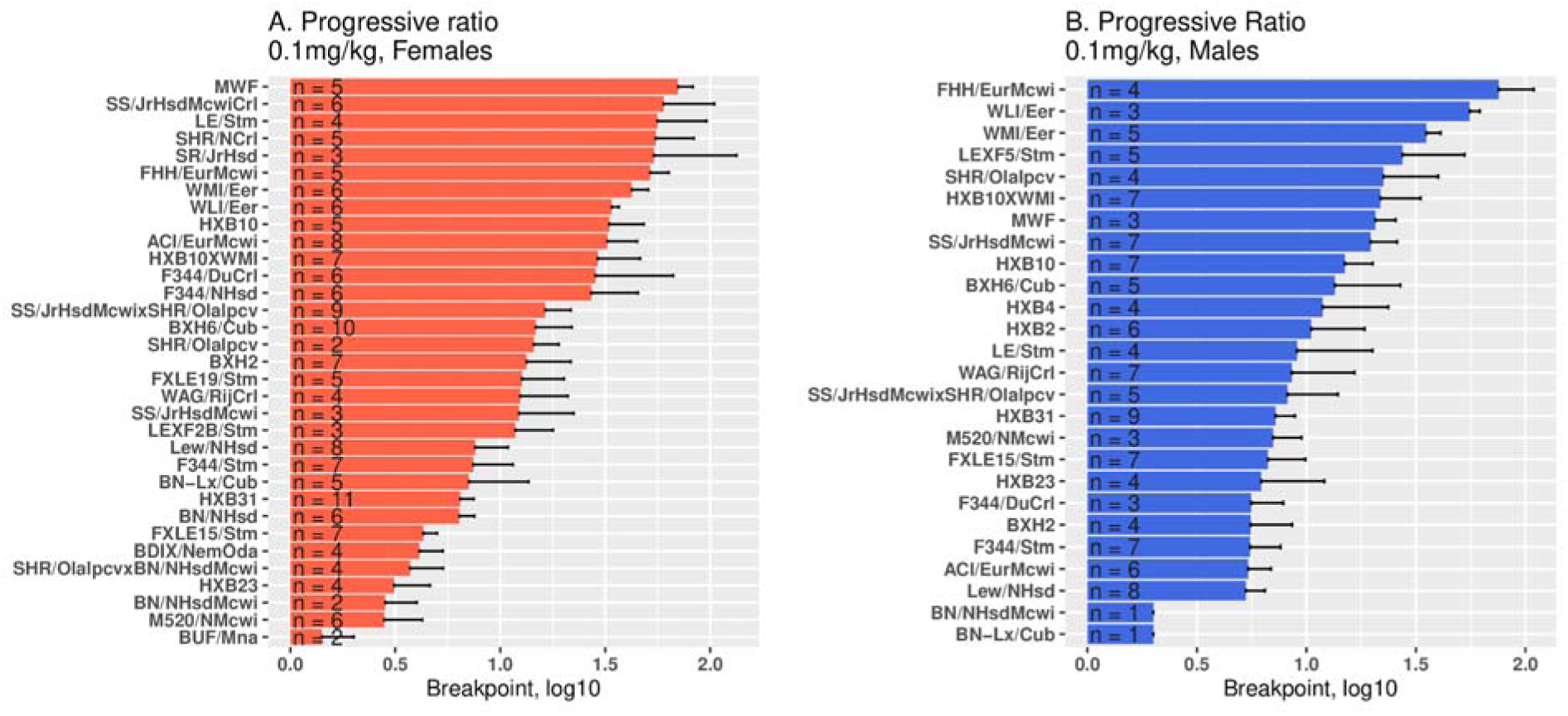
Breakpoints during progressive ratio schedule of increasing active licks per oxycodone dose (0.1 mg/kg). In each sex, the breakpoints reached were significantly strain-dependent (A. females, F_32,148_=5.2, p=1.8e-12; B. males, F_25,103_=2.6, p=0.0003) and greater in females than males (F_22,227_=6.4, p=2.5e-14). X-axis is in log scale.

During cue-induced reinstatement of extinguished oxycodone seeking (Figure 8), we found a strain difference throughout the HRDP in the number of active licks in females (A. F_31,142_=2.9, p=1.3e-5) and males (B. F_25,94_=3.2, p=2.0e-5). In the 23 strains where reinstatement was studied in both sexes, female active licks were greater (F_23,221_=3.8, p=1.5e-7). Hence, cue-induced oxycodone-seeking is strain-dependent and stronger in females than males.

**Figure 8.**
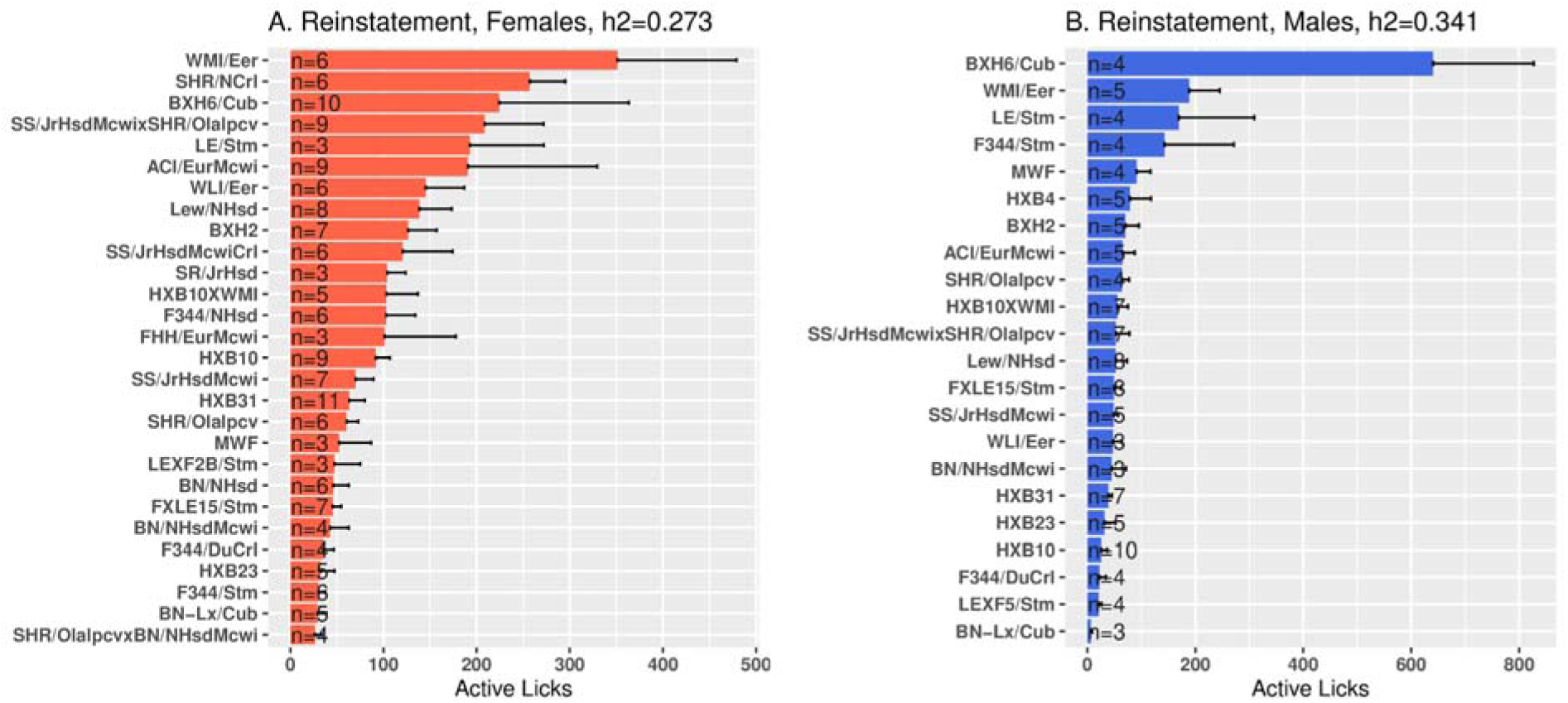
Cue-induced reinstatement of extinguished oxycodone seeking. Reinstatement was conducted after the number of licks on the active spout was less than 100 during two consecutive extinction sessions. There was a strain difference in the number of active licks during reinstatement in females (A. F_31,142_=2.9, p=1.3e-5) and males (B. F_25,94_=3.2, p=2.0e-5). In the 23 strains where reinstatement was studied in both sexes, female active licks were greater (F_23,221_=3.8, p=1.5e-7).

### The Correlation between Male (M) vs. Female (F) Oxycodone Intake

In the 23 common strains, we found inter-sex correlations in oxycodone (0.1 mg/kg) intake during 4-h and 16-h sessions. During 4-h sessions with stable oxycodone intake, male vs. female intake was highly correlated across strains (Figure 8A, Pearson r=0.569, p=0.005). However, during 16-h sessions, intake between sexes was not correlated across the HRDP (Figure 8B, r=0.374, p=0.104). The heritability of oxycodone intake declined in males from 4-h to 16-h sessions (Table 2; *h*^2^ 0.633 to 0.222) and this also occurred, to some extent, in females (*h*^2^ 0.383 to 0.299); this decrease indicated increased intra-strain variability in intake during the extended access sessions, which could reduce the correlation between sexes (Figure 8B).

**Table 2.**
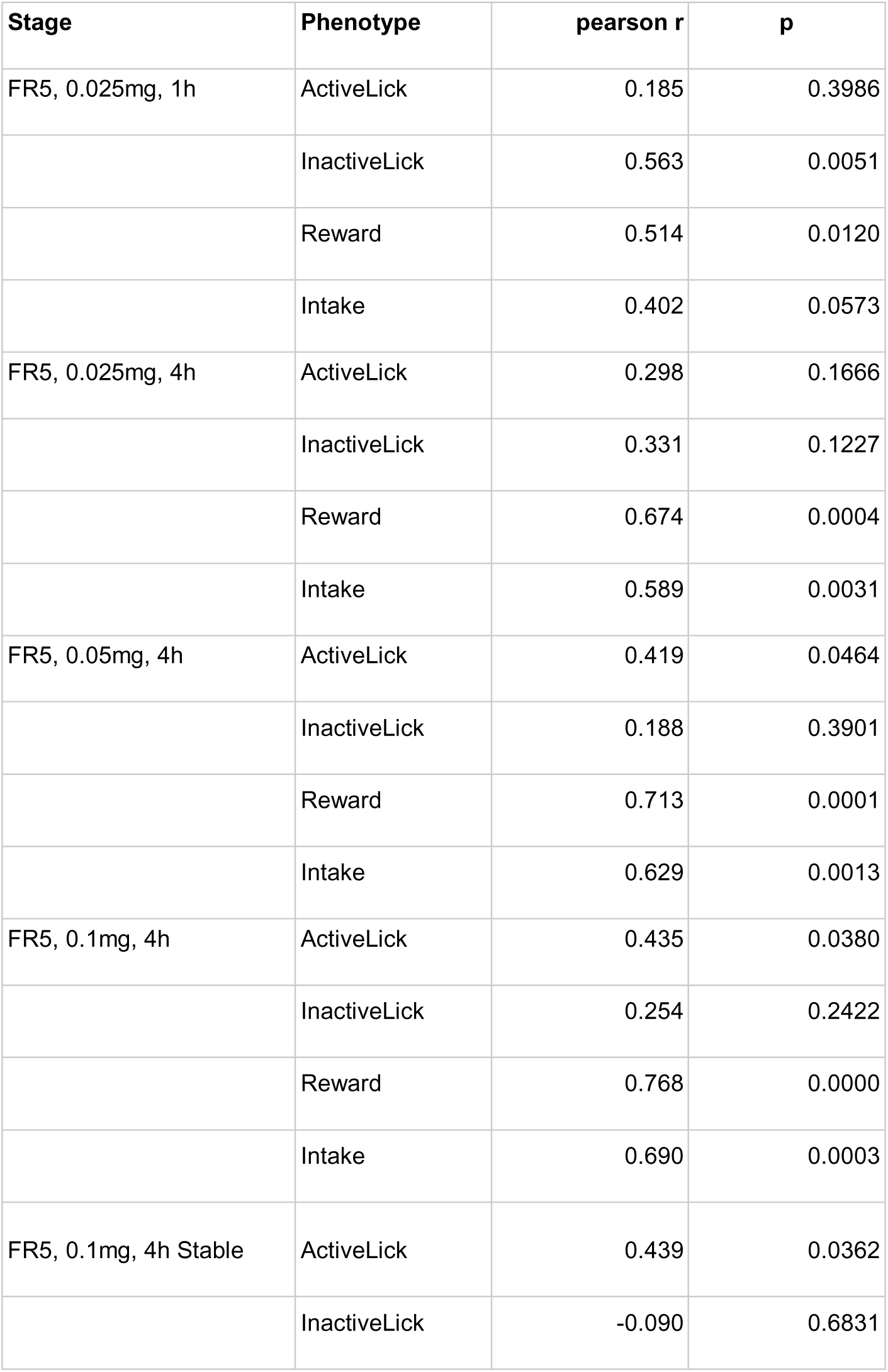
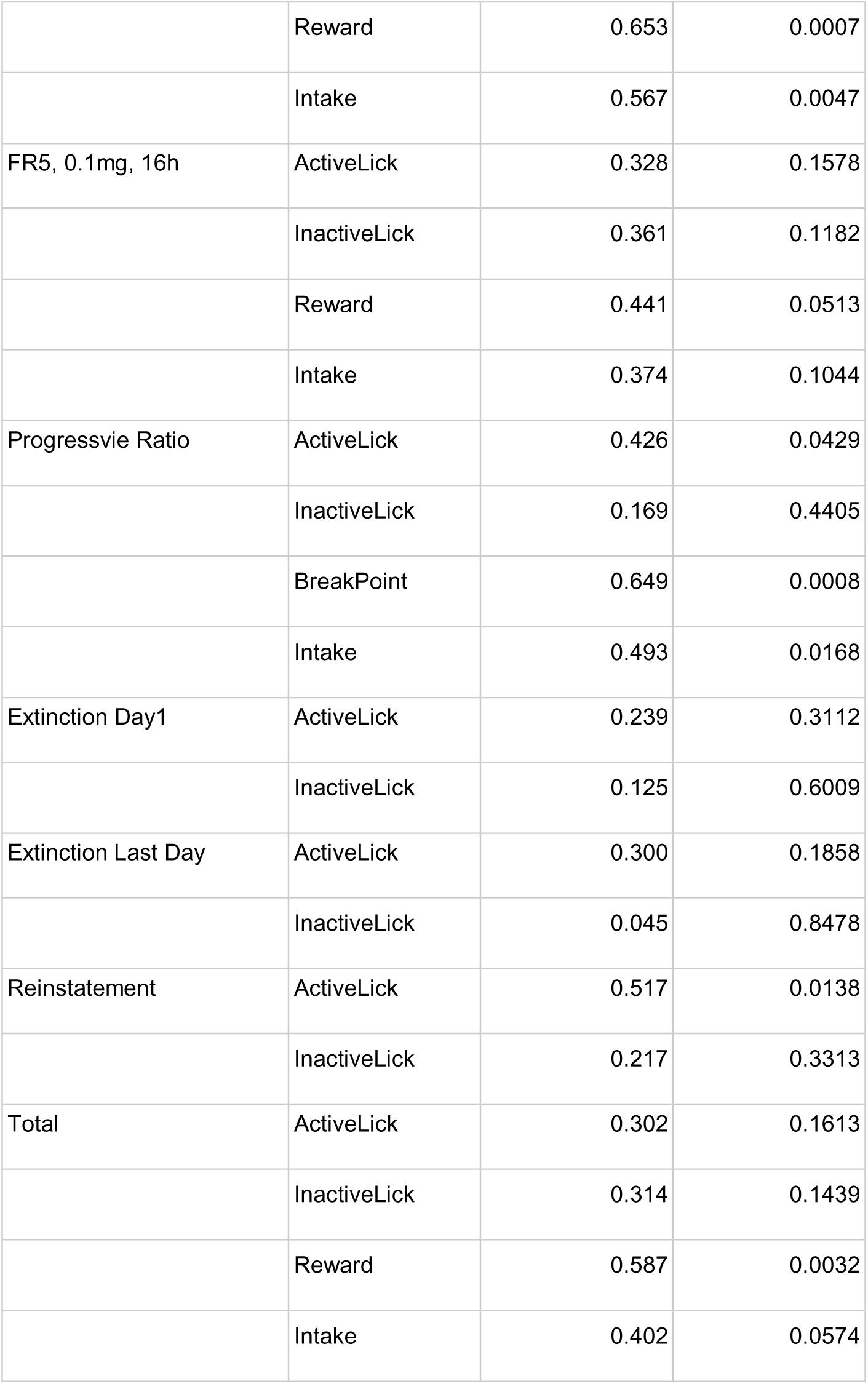
In the 23 common strains, the correlation between males and females in oxycodone self-administration.

In Figure 9A, 4-h intake was highly correlated between sexes across the 23 strains, despite strong sex differences in oxycodone intake (Figure 3), motivation to obtain drug (Figure 7), and seeking behavior during reinstatement (Figure 8). This demonstrates the strong strain-dependent genetic control of oxycodone intake that co-exists with sex-dependent behaviors. Table 2 underscores this strong strain-dependent genetic control: multiple parameters of oxycodone self-administration were significantly correlated between sex across the 23 strains These are: oxycodone intake during 1-h, 4-h and 16-h sessions, effort to obtain reward (i.e., breakpoint and rewards obtained during PR trial), and active licks during reinstatement.

**Figure 9.**
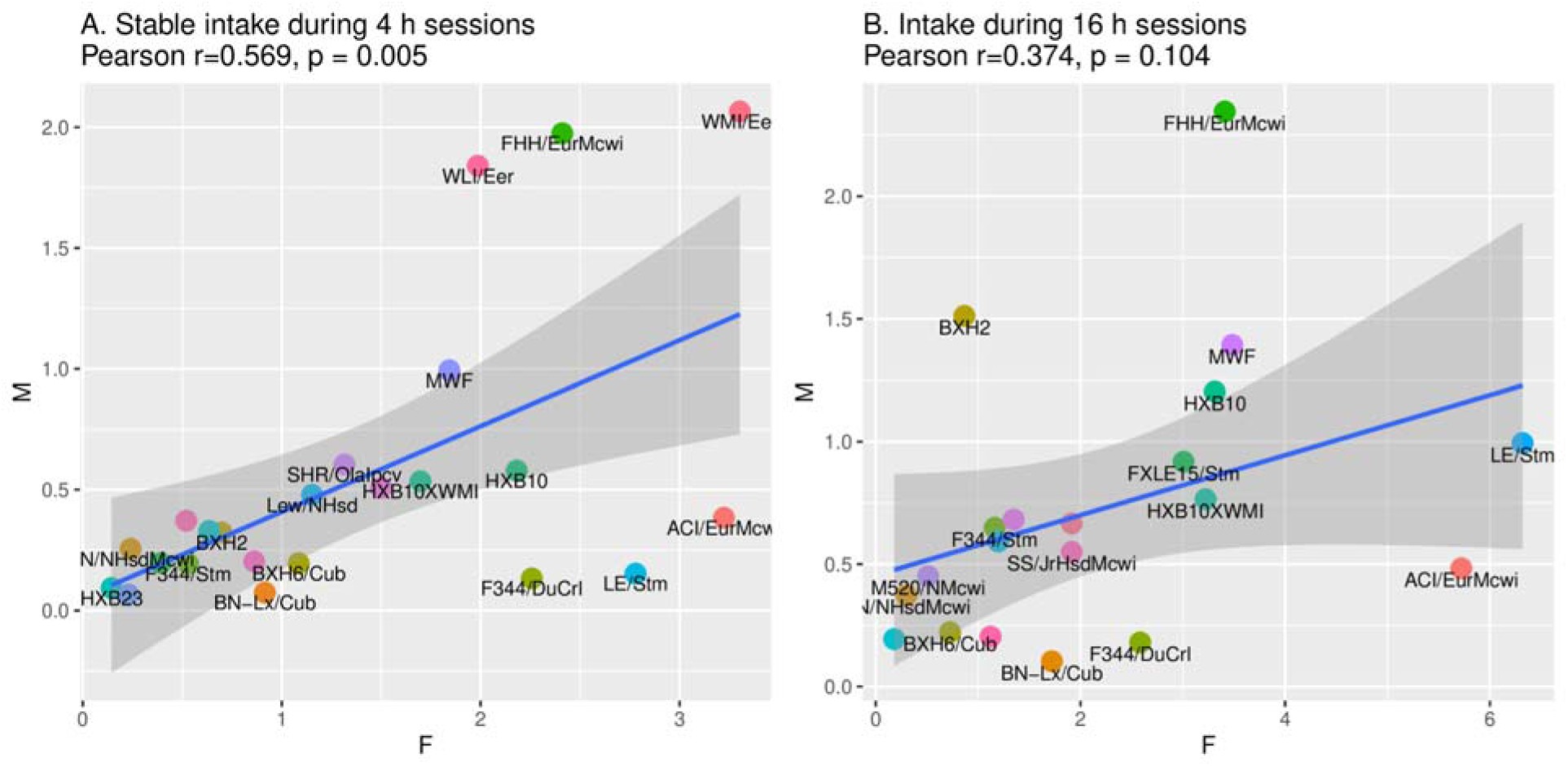
In the 23 common strains, the correlation in male (M) vs. female (F) oxycodone (0.1 mg/kg) intake/session during 4-h and 16-h sessions. A. During 4-h sessions with stable oxycodone intake, M vs F intake was highly correlated across the HRDP (Pearson r=0.54, p=0.008). B. However, during 16-h sessions, intake between sexes was not significantly correlated (r=0.347, p=0.104).

### The Heritability (*h*^2^) of Oxycodone Behavioral Phenotypes

Table 3 shows the heritability (*h*^2^) of oxycodone SA parameters by sex. Active licks, reward, and intake show h^2^ values associated with significant differences between strains for all SA parameters in each sex. Inactive licks were generally at much lower h^2^ values in each sex. In males, the *h*^2^ for 4-h oxycodone intake (mg/kg b.wt./session) was consistently greater than in females by approximately 50%. In contrast, *h*^2^ for active licks in 4-h sessions was higher in females. In 16-h sessions, the *h*^2^ for both intake and active licks was greater in females than males. Additionally, the female *h*^2^ for intake under a. progressive ratio schedule was 0.41 vs. 0.31 in males; active licks under this schedule showed a greater divergence in *h*^2^ between female and male: 0.46 and 0.15, respectively.

**Table 3.** Heritability and ANOVA results for oxycodone SA parameters.

|  | Phenotype | Males |  |  |  | Females |  |  |  |
| --- | --- | --- | --- | --- | --- | --- | --- | --- | --- |
|  |  | h2 | Df | F | p | h2 | Df | F | p |
| FR5, 0.025mg,<br>1h | Active Lick | 0.302 | (25,441) | 7.77 | 1.3E-22 | 0.368 | (32,588) | 10.87 | 2.2E-41 |
|  | Inactive Lick | 0.216 | (25,436) | 5.20 | 8.4E-14 | 0.227 | (32,579) | 5.89 | 3.0E-20 |
|  | Reward | 0.341 | (25,441) | 9.13 | 4.9E-27 | 0.261 | (32,588) | 6.99 | 4.0E-25 |
|  | Intake | 0.347 | (25,432) | 9.07 | 9.5E-27 | 0.287 | (32,570) | 7.45 | 5.3E-27 |
| FR5, 0.025mg,<br>4h | Active Lick | 0.297 | (25,277) | 5.34 | 3.3E-13 | 0.411 | (32,376) | 8.71 | 2.9E-29 |
|  | Inactive Lick | 0.081 | (25,272) | 1.86 | 9.0E-03 | 0.252 | (32,366) | 4.63 | 1.2E-13 |
|  | Reward | 0.443 | (25,277) | 9.15 | 3.8E-24 | 0.342 | (32,376) | 6.75 | 5.2E-22 |
|  | Intake | 0.439 | (25,275) | 8.95 | 1.4E-23 | 0.297 | (32,374) | 5.65 | 9.8E-18 |
| FR5, 0.05mg, 4h | Active Lick | 0.298 | (25,276) | 5.31 | 4.2E-13 | 0.469 | (32,374) | 10.82 | 1.9E-36 |
|  | Inactive Lick | 0.156 | (25,276) | 2.84 | 1.6E-05 | 0.228 | (32,358) | 4.11 | 1.5E-11 |
|  | Reward | 0.655 | (25,278) | 20.38 | 5.2E-49 | 0.417 | (32,374) | 8.95 | 4.5E-30 |
|  | Intake | 0.652 | (25,278) | 20.12 | 1.6E-48 | 0.406 | (32,374) | 8.59 | 9.0E-29 |
| FR5, 0.1mg, 4h | Active Lick | 0.295 | (25,277) | 5.23 | 7.0E-13 | 0.531 | (32,372) | 13.40 | 1.7E-44 |
|  | Inactive Lick | 0.072 | (25,275) | 1.78 | 1.4E-02 | 0.158 | (32,363) | 2.96 | 4.5E-07 |
|  | Reward | 0.663 | (25,277) | 20.93 | 6.3E-50 | 0.378 | (32,372) | 7.65 | 2.5E-25 |
|  | Intake | 0.654 | (25,277) | 20.17 | 1.5E-48 | 0.367 | (32,372) | 7.36 | 2.8E-24 |
| FR5, 0.1mg, 4h<br>Stable | Active Lick | 0.39 | (25,506) | 10.17 | 2.9E-31 | 0.509 | (32,620) | 15.33 | 4.4E-59 |
|  | Inactive Lick | 0.206 | (25,505) | 4.65 | 4.2E-12 | 0.252 | (32,615) | 5.53 | 7.9E-19 |
|  | Reward | 0.644 | (25,506) | 26.86 | 1.1E-76 | 0.369 | (32,620) | 9.10 | 1.5E-34 |
|  | Intake | 0.633 | (25,506) | 25.74 | 3.8E-74 | 0.383 | (32,620) | 9.59 | 1.3E-36 |
| FR5, 0.1mg, 16h | Active Lick | 0.25 | (22,352) | 5.60 | 2.0E-13 | 0.396 | (25,426) | 10.92 | 2.1E-32 |
|  | Inactive Lick | 0.149 | (22,350) | 3.41 | 7.0E-07 | 0.122 | (25,425) | 3.10 | 1.4E-06 |
|  | Reward | 0.252 | (22,352) | 5.65 | 1.4E-13 | 0.292 | (25,426) | 7.25 | 1.0E-20 |
|  | Intake | 0.222 | (22,352) | 4.94 | 1.9E-11 | 0.299 | (25,426) | 7.44 | 2.3E-21 |
| Progressvie<br>Ratio | Active Lick | 0.147 | (23,103) | 1.82 | 2.2E-02 | 0.458 | (29,145) | 5.42 | 2.7E-12 |
|  | Inactive Lick | 0.083 | (23,101) | 1.41 | 1.2E-01 | 0.212 | (29,145) | 2.41 | 3.4E-04 |
|  | BreakPoint | 0.256 | (23,103) | 2.64 | 4.6E-04 | 0.44 | (29,145) | 5.10 | 1.7E-11 |
|  | Intake | 0.311 | (23,103) | 3.14 | 3.8E-05 | 0.408 | (29,145) | 4.60 | 3.6E-10 |
| Extinction Day1 | Active Lick | 0.316 | (21,99) | 3.24 | 4.4E-05 | 0.299 | (25,125) | 3.20 | 1.0E-05 |
|  | Inactive Lick | 0.142 | (21,99) | 1.80 | 2.9E-02 | 0.106 | (25,125) | 1.61 | 4.6E-02 |
| Extinction Last<br>Day | Active Lick | 0.22 | (20,81) | 2.26 | 5.4E-03 | 0.275 | (25,127) | 2.95 | 3.9E-05 |
|  | Inactive Lick | 0.036 | (20,81) | 1.17 | 3.1E-01 | 0.221 | (25,127) | 2.46 | 5.9E-04 |
| Reinstatement | Active Lick | 0.341 | (21,92) | 3.41 | 2.5E-05 | 0.273 | (27,140) | 2.94 | 2.0E-05 |
|  | Inactive Lick | 0.07 | (21,92) | 1.35 | 1.7E-01 | 0.255 | (27,140) | 2.76 | 5.9E-05 |
| Total | Active Lick | 0.217 | (25,2395) | 22.84 | 7.3E-93 | 0.305 | (32,3097) | 37.86 | 3.0E-195 |
|  | Inactive Lick | 0.122 | (25,2380) | 11.63 | 4.5E-44 | 0.142 | (32,3047) | 14.40 | 2.0E-71 |
|  | Reward | 0.326 | (25,2397) | 39.14 | 8.6E-158 | 0.181 | (32,3097) | 19.50 | 5.1E-100 |
|  | Intake | 0.196 | (25,2386) | 20.14 | 2.4E-81 | 0.146 | (32,3077) | 15.21 | 4.8E-76 |

Overall, the relative degree of oxycodone heritability in males vs. females was specific for each behavioral phenotype. Oxycodone intake in 4-h sessions at three doses showed the highest heritability and some of the largest differences between sexes, with males considerably greater than females. However, active licks in 4-h sessions and both intake and active licks in 16-h sessions and under a PR schedule were greater in females than males.

### Correlations between Behavioral Tests and Oxycodone SA

We assessed anxiety-like and response to novelty using elevated plus maze (EPM), open field test (OFT) and novel object interaction (NOI) in *naive* rats. Table 4 shows that the significance of the traits defining each behavioral test (i.e., 5 EPM traits, 4 OFT, 4 NOI) varied within each sex across the HRDP strains we tested (n=19 female, 17 male). Additionally, Table 4 shows that the greatest *h*^2^ values within sex were found for total distance in all three behavioral tests (range across tests by sex: female, 0.51-0.58; male, 0.24-0.49).

**Table 4.** Heritability and ANOVA for comparison across HRDP of anxiety-like and novelty response traits in naive inbred rats.

| Behavior | Trait | Males |  |  |  | Females |  |  |  |
| --- | --- | --- | --- | --- | --- | --- | --- | --- | --- |
|  |  | h2 | Df | F | p | h2 | Df | F | p |
| OFT | totalDistance | 0.493 | (15,96) | 6.41 | 3.19E-09 | 0.585 | (15,105) | 10.06 | 2.19E-14 |
| OFT | dist2CtrMean | 0.553 | (15,96) | 7.87 | 3.26E-11 | 0.437 | (15,105) | 5.98 | 7.96E-09 |
| OFT | cntrDurat | 0.326 | (15,96) | 3.69 | 4.44E-05 | 0.123 | (15,105) | 1.90 | 3.07E-02 |
| OFT | cntrFreq | 0.334 | (15,96) | 3.79 | 3.09E-05 | 0.395 | (15,105) | 5.19 | 1.32E-07 |
| NOI | totalDistance | 0.398 | (15,100)<br>) | 4.89 | 4.74E-07 | 0.562 | (16,114) | 9.06 | 5.71E-14 |
| NOI | dist2CtrMean | 0.276 | (15,100)<br>) | 3.25 | 2.13E-04 | 0.248 | (16,114) | 3.07 | 2.54E-04 |
| NOI | cntrDurat | 0.105 | (15,100)<br>) | 1.69 | 6.40E-02 | 0.152 | (16,114) | 2.13 | 1.15E-02 |
| NOI | cntrFreq | 0.272 | (15,100)<br>) | 3.20 | 2.58E-04 | 0.489 | (16,114) | 7.03 | 5.35E-11 |
| EPM | totalDistance | 0.237 | (16,88) | 2.75 | 1.31E-03 | 0.510 | (18,101) | 6.69 | 1.12E-10 |
| EPM | openDurat | 0.191 | (16,88) | 2.34 | 6.27E-03 | 0.216 | (18,101) | 2.50 | 2.04E-03 |
| EPM | openFreq | 0.295 | (16,88) | 3.37 | 1.29E-04 | 0.241 | (18,101) | 2.74 | 7.68E-04 |
| EPM | closedDurat | 0.300 | (16,88) | 3.42 | 1.08E-04 | 0.118 | (18,101) | 1.73 | 4.60E-02 |
| EPM | closedFreq | 0.225 | (16,88) | 2.64 | 2.01E-03 | 0.339 | (18,101) | 3.80 | 8.58E-06 |

We correlated multiple traits measured in each behavioral paradigm with 28 parameters of oxycodone SA, using the complete HRDP dataset. We used this dataset to find the maximum number of significant correlations within each sex with the proviso that deductions should not be drawn from inter-sex comparisons of the number of significant correlations. Figure 10 shows examples of two significant correlations for EPM within sex vs. 4-h intake of oxycodone 0.025 (panels A, B) and 0.05 (panels E, F). Figure 11 shows that EPM-defining traits (i.e., female dataset, 13; male dataset, 7) were associated with multiple SA parameters in each sex, and a large fraction of these associations were sex specific (e.g., female only 7/13). In females, three EPM traits (Figure 11) correlated with total 4-h oxycodone intake and one trait with 4-h intake in males. OFT traits were SA-associated in each sex: 8 in males and 2 in females, and specific SA associations were unique to each sex (Figure 11). NOI traits were associated with four SA parameters in males and none in females. Amongst these male NOI associations, NOI correlated with total oxycodone intake (i.e., sum of all doses at all time intervals in males/strain) and with active licks during reinstatement compared to OFT and EPM in reinstating females. Figure 10D shows the strain correlation for NOI (distance to center) and total oxycodone intake. At the highest dose of oxycodone (0.1 mg/kg), no behavioral traits correlated with intake, reward or active licks at 4-h and 16-h, barring one exception - active licks in males during 4-h sessions correlated with EPM. This is the same oxycodone dose consumed in the PR study that correlated with NOI and OF only in males.

**Figure 10.**
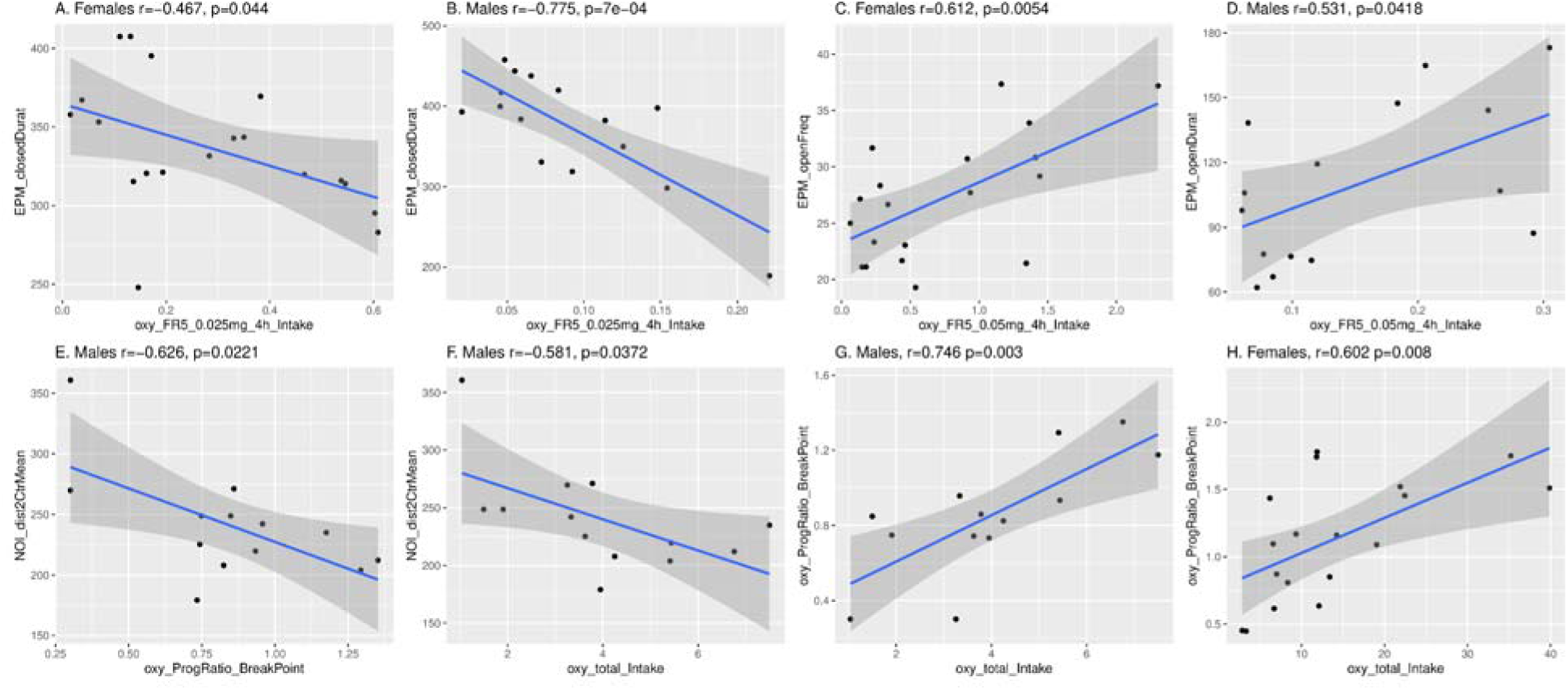
Representative correlations between oxycodone SA parameters and behavioral traits. Panels A-D show significant correlations for EPM vs. oxycodone intake in 4-h SA sessions at 0.25 and 0.5 mg/ml in female (panels A, C) and male (B, D) HRDP strains. Significant correlations in male HRDP strains between NOI (distance to center) and PR (breakpoint) or total oxycodone intake are in panels E and F, respectively. Correlations between NOI and PR or total intake were not significant in females (see supplementary Figure S1). Panels G and H, show significant correlations between two SA parameters, PR and total intake, in males and females, respectively.

**Figure 11.**
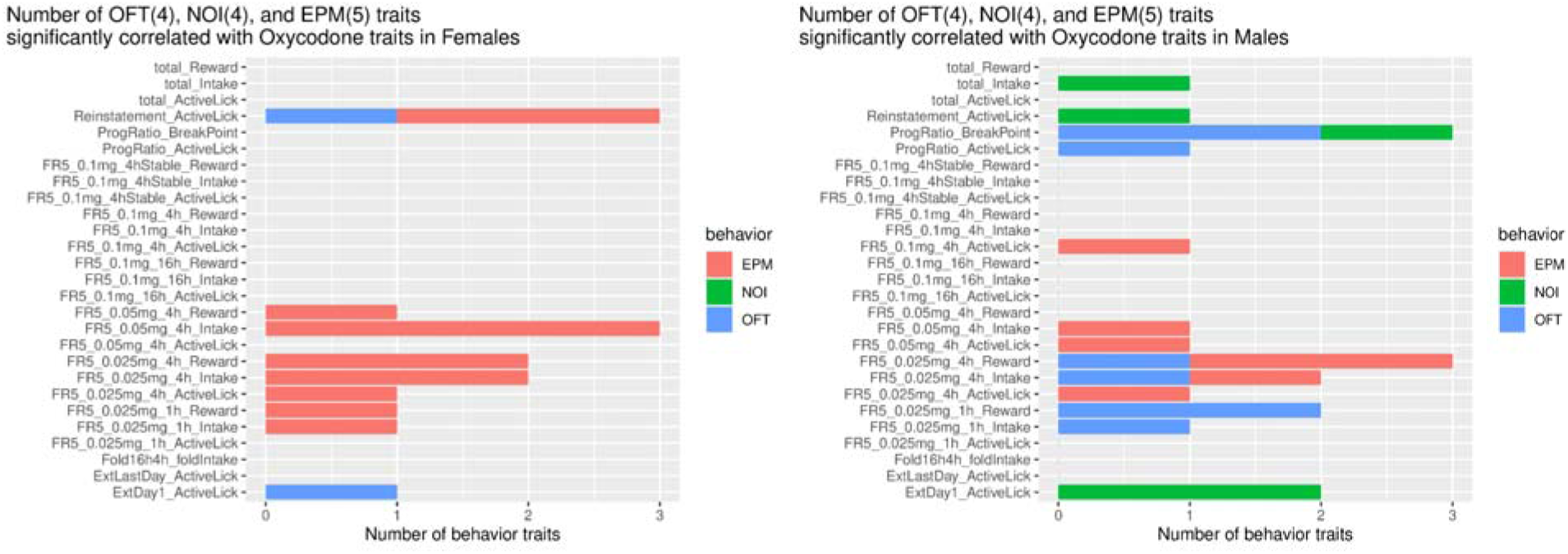
Behavioral traits that significantly correlate with oxycodone SA parameters. This bar graph groups all the behavioral traits measured in EPM, OFT, and NOI trials that were associated with each SA parameter; this grouping is based on the significant correlations (p<0.05) identified between a single behavioral trait and an SA parameter (supplementary data, figures S1 and S2).

### Correlation in Males between PR Breakpoint, NOI (distance to center), and Total Oxycodone Intake

NOI (distance to center) correlated with both PR breakpoint and total oxycodone intake in males, but not females (Figure 11); indeed, no behavioral traits were associated with either of these two SA parameters in females. Figure 10C shows this correlation for NOI and PR breakpoint in males. In Figure 10 (panels G, H), PR breakpoint also significantly correlated across strains with total oxycodone intake in both sexes. In summary, the following correlations were identified in males across strains: PR breakpoint x NOI (distance to center); total oxycodone intake x NOI; PR breakpoint x total oxycodone intake. Hence, these three independent measures are all significantly inter-correlated across strains only in males. This strongly suggests that CNS mechanisms governing novelty seeking and motivation to take oxycodone interact in males to regulate the total consumption of oxycodone.

## Discussion

Human inter-individual variation in the susceptibility to drug abuse is characteristic of addiction to opioids. Outbred animal models also demonstrate substantial inter-individual variation in responses to behavioral paradigms with face validity for important dimensions of human addiction ^14^. We took advantage of the highly replicable behaviors amongst individuals within inbred rat strains to identify behavioral parameters of oxycodone SA that consistently varied across a large panel of inbred strains (i.e. hybrid rat diversity panel, HRDP) - bred, raised and tested in the same vivarium. Under these conditions, consistent inter-strain variation in oxycodone SA should reflect differences in the genetic regulation of CNS functions controlling SA parameters. Indeed, we identified multiple heritable parameters of oxycodone SA that also showed sex-dependency. The broad polygenic regulation of each behavioral parameter and the impact of subsets of genes on multiple parameters implies a strong functional inter-dependency amongst these parameters; hence, they do not operate as independent variables. Many of these SA parameters correlated with the responses in naive HRDP rats to independent behavioral testing in EPM, NOI and OFT.

Oxycodone intake, both during initiation of SA and stable intake in 4-h sessions, was strain-dependent in both sexes (Figure 4) and heritable (Table 3). In the 23 strains common to both sexes, the mean amount consumed across all sessions (Figure 3) of increasing duration and dose was greater in females than males (p<0.001), and the mean number of active licks (Figure 2) was greater in females during 4-h (p=0.003) and 16-h sessions (p=0.03)). Similar to 4-h sessions, oxycodone intake in 16-h sessions was heritable in both sexes (Table 2). Within sex, oxycodone intake was correlated in 4-h vs. 16-h sessions (Figure 5C, 5D). Therefore, in individual female and male strains, intake in 4-h sessions predicted intake during extended access sessions. This within-strain correlation of oxycodone intake in 4-h and 16-h sessions is most probably controlled by the heritability of genes that regulate intake in both short and extended access sessions.

Oxycodone intake by a subset of strains in each sex was dramatically augmented (>3-fold) during 16-h sessions. Augmented intake was evident within the first 6-h and rapidly declined thereafter (Figure 6). Considering the half-life of blood oxycodone in rat ^15^, it is likely that an approximate doubling of blood levels occurred during this interval while intake and blood levels declined rapidly in non-augmenters. This suggests that oxycodone is either more rewarding in augmenters or that more drug is required to produce adequate reward, perhaps due to lower sensitivity or higher clearance. Since hourly and total intake in 4-h sessions were similar between augmenters and non-augmenters in both sexes (Figure 6), reduced sensitivity to the rewarding effects is unlikely in augmenters. Similarly, enzymatic clearance of blood oxycodone is unlikely to vary between the two groups. Hence, we suggest that oxycodone is probably more rewarding in augmenters due to the genetic control of reward efficacy.

In general, females across all strains manifest greater oxycodone intake in both short and extended access sessions and greater numbers of active licks. This strongly suggests the existence of a basic sex difference in the amount of oxycodone required to establish stable oral oxycodone reinforcement within the CNS. An extensive literature on sex differences in the sensitivity to opiate reinforcement and analgesia supports this hypothesis.

Morphine has been reported to induce a more pronounced place preference in female Wistar rats at similar doses ^16^. In self-administering female Sprague Dawley rats, more intravenous (i.v.) morphine and heroin were consumed and a broader range of doses were reinforcing than in males ^17^. Moreover, at doses in the upper end of the dose-response range, morphine-induced place preference in females, but was no longer effective in males ^18^. Both estrogen and progesterone receptors have been detected in dopamine terminals and median spiny neurons within nucleus accumbens (NAc), which is involved in reward-associated learning and motivation to goal-oriented behaviors ^19,20^. Estradiol rapidly enhanced NAc dopamine release and modulated dopamine binding, effects also observed in cycling females ^19^ In mice, basal dopamine neuron activity and dopamine release in NAc were similar in both sexes, except during estrus when both increased ^21^. Additionally, both systemic and intrastriatal estradiol rapidly amplified amphetamine-induced dopamine release in rat dorsal striatum ^22^. In summary, rat models of opiate preference and operant self-administration demonstrate that opiates are more reinforcing in females, and at a broader and higher dose range. This is likely due, in part, to enhanced responsiveness to opiate-induced dopamine release under the influence of ovarian steroids. Hence, oral oxycodone appears to be more reinforcing in females across the HRDP, which drives greater licking behavior.

PR breakpoints were higher in female strains across the HRDP. Similar differences have been reported for i.v. opiate SA in outbred rats ^17^. Since PR breakpoint is a heritable SA parameter in both sexes, it is reasonable to expect significant inter-strain variability in the effect of sex if the overall effect of sex is small to moderate and the interaction between sex and heredity in each strain depends on the specific subsets of genes modulating PR in each strain.

Multiple traits measured during independent behavioral tests, conducted in naive rats, varied significantly across strains and were heritable (Table 4). These traits correlated with specific parameters of oxycodone SA, depending on sex (Figure 10 and supplementary Figures S1, S2). EPM traits were associated with a common subset of SA parameters in both sexes, whereas NOI traits were associated with SA only in males. In both 4-h and 16-h sessions at high dose oxycodone, with one exception (Figure 10: male, EPM vs. active licks, 4-h), no SA parameters correlated with behavior. At high dose oxycodone, intake and active licks in short and long access sessions were not correlated with behavioral measures of anxiety (i.e., EPM and OFT) and novelty-seeking (i.e., NOI). Hence, the reinforcing efficacy of high dose oxycodone is unaffected by the intrinsic, strain-dependent level of anxiety or novelty-seeking associated with oxycodone intake at lower doses.

The NOI trait of distance to the center was inversely associated with both PR breakpoint and total oxycodone intake in males (Figure 9, 10). In contrast, no correlations were found between these SA parameters and behavioral traits in females (Figure 11). Overall, the heritable, sex-specific correlation of SA parameters with specific behavioral traits indicates the influence of pleiotropic genes affecting both parameters of oxycodone SA and specific behavioral traits modulated by sex, Significant 3-way correlations of PR breakpoint x NOI-distance to center, PR breakpoint x total oxycodone intake, and total oxycodone intake x NOI-distance to center strongly suggest that one set of pleiotropic genes may underlie these correlations in males. It is most likely that these genes directly modulate oxycodone and novelty seeking behavior, because behavioral tests were conducted in drug naive individuals. Novelty-seeking and motivation to take oxycodone may interact and regulate oxycodone intake depending on the strain-specific expression of these pleiotropic genes in males.

These studies demonstrate the strong strain-dependent inheritance of multiple oxycodone SA parameters and their correlations with independent measures of anxiety and novelty in established behavioral tests (i.e., EPM, OFT, NOI). Overall, active licks and oxycodone intake in multiple phases of our experimental protocol were significantly greater in females of most strains. In both sexes, 16-h vs 4-h intake increased and 4-h intake predicted 16-h intake in both sexes. A subset of strains in each sex dramatically augmented their intake during the first 6-h of 16-h sessions, suggesting genetic regulation of reward efficacy. The genome-wide search for quantitative trait loci (QTLs) based on mapping these strongly heritable SA parameters and behavioral traits is likely to yield positive results, especially when these parameters and traits (e.g., PR-breakpoint, total oxycodone intake, NOI-distance to center) involve pleiotropic genes. Such genes increase the confidence of finding strong gene candidates in QTLs because they are likely to be found due to overlapping single nucleotide polymorphisms (SNPs) discovered by independently mapping of several different behaviors.

## Methods

### Animals

Breeders from the HRDP were obtained from Dr. Melinda R. Dwinell at the Medical College of Wisconsin. All rats were bred on campus and housed in groups in a room with a 12:12 h reversed light cycle at the University of Tennessee Health Science Center. Experiments were conducted during the dark phase of this cycle. Each rat, including breeders and offspring, had a radio frequency identification (RFID) tag implanted under its skin for identification purposes. Adult rats (PND 65-90) of both sexes participated in oxycodone self-administration experiments. A separate group of adolescent rats (PND38-44) underwent behavioral tests. The Animal Care and Use Committee of The University of Tennessee Health Science Center approved all procedures, which complied with NIH Guidelines for the Care and Use of Laboratory Animals.

### Drugs

Oxycodone HCl was a kind gift by Noramco (Wilmington, DE).

### Open Field Test

OFT was carried out as previously reported ^23^. Two OFT arenas were constructed using black acrylic glass, measuring 100 cm (L) × 100 cm (W) × 50 cm (H), which were placed side by side. The floors were covered by wood boards painted with either black or white acrylic paint (ART-Alternatives, ASTM D-4236, Emeryville, CA, USA) to contrast the coat of the animals (i.e., a black board was used for rats with white fur). The test chambers were illuminated by a long-range, 850-nm infrared light (LIR850-70, LDP LLC, Carlstadt, NJ) located 160 cm above the center of the two test chambers. No source of visible light was present during behavioral testing, with the exception of a flat panel monitor (Dell 1908FP). A digital camera (Panasonic WV-BP334) fitted with an 830 nm infrared filter (X-Nite830-M37, LTP LLC, Carlstadt, NJ) and located next to the infrared light source was used to record the behavior of the rats. All rats were released at the same corner of the test chamber, and data were collected for 20 min.

### Novel Object Interaction (NOI) Test

This test was conducted the day after the OFT in the same arena. A cylindrical rat cage constructed using 24 aluminum rods (30 cm in length) spaced 1.7 cm apart was used as the novel object ^24^. The bottom and top of the cage (15 cm in diameter) were manufactured using a 3D printer from polylactic acid. The novel object was placed in the center of the arena before testing. The test duration was 20 min and was recorded using the same camera as that used in the OFT.

### Elevated Plus Maze

The maze was constructed using black acrylic glass. The floors of the maze were covered by wood boards painted with black or white acrylic paint. The platform was 60 cm above the floor, with all four arms measuring 12 cm (W)L×L50Lcm (L). The two opposing closed arms had walls measuring 30 cm (H). Rats were placed into the center of the maze facing the closed arm. The behavior of the rat was recorded for 6 min using the digital video system described above.

### Analysis of Video Data

Ethovision XT video tracking system (RRID:SCR_000441, Version 15.0, Noldus Information Technology, The Netherlands) was used to analyze the videos recorded in all behavioral tests. After identifying the arena and calibrating the size of the arena, specific zones in the arena were outlined. For OFT and NOI, one center zone, which was a circular region with a diameter of 20 cm, was used. The extracted data included the total distance traveled in the arena, the duration and the frequency the test rat was present in specific zones, and the distance of the subject to the zones. The center of the subject rat was used for all calculations.

### Oral Operant Oxycodone Self-Administration

We employed the same methodology as previously reported ^8^, with minor adjustments. The operant chamber (Med Associates) featured two lickometers: licks on the active spout following a fixed ratio 5 (FR5) schedule triggered the immediate release of a 60 μl oxycodone solution (0.025–0.1 mg/ml) onto the spout’s tip, along with the activation of an LED visual cue. A 20-second timeout followed the drug delivery, during which licks on the active spout and any licks on the inactive spout were recorded but had no programmed consequences. Rats had free access to food and water and were kept under a reversed light cycle, being tested in their dark phase.

Training commenced with three daily 1-h FR5 sessions at 0.025 mg/ml oxycodone concentration. Subsequent sessions extended to 4-h and occurred every other day. From session four onwards, we doubled the dose every two sessions up to the maximum dose of 0.1 mg/ml. Rats underwent six sessions at this highest dose, followed by a progressive ratio test on session fourteen. Session lengths then increased to 16-h (4 PM - 8 AM) for three sessions. Extinction sessions were conducted for 1-h daily, without programmed consequences, until licks on the active spout decreased to less than fifty for two consecutive sessions. A final reinstatement session was carried out where active spout licks triggered only visual cue delivery without oxycodone. We recorded both the number and timing of oxycodone deliveries as well as licks on active and inactive spouts. The procedure is summarized in Figure 1. Full strain names and sexes of rats involved in oxycodone self-administration, along with the number of rats per strain (minimum is 3) and sex are listed in Table 1.

### Identification of augmenter vs non-augmenter strains

In each sex, HRDP strains with more than a threefold increase in oxycodone (0.1 mg/ml) intake during 16-h vs 4-h sessions were classified as augmenters: 7 females, FXLE19/Stm, WAG/RijCrl, SR/JrHsd, LEXF2B/Stm, SHR/OlaIpcvxBN/NHsdMcwi, FXLE15/Stm, BDIX/NemOda; 6 males, F344/Stm, LEXF5/Stm, FXLE15/Stm, BXH2, LE/Stm, M520/NMcwi). In each sex, all HRDP strains were then rank ordered by intake in 4-h sessions. For each augmenter strain, one strain that ranked immediately above it (i.e., higher 4-h intake) and not already identified as an augmenter or non-augmenter was specified as the control non-augmenter strain. These are: females, SHR/OlaIpcv, BXH2, BN/NHsdMcwi, HXB23, HXB31, F344/Stm, M520/NMcw; males, SS/JrHsdMcwixSHR/Olalpcv, BXH6/Cub, F344/DuCrl, BN-Lx/Cub, HXB31, HXB23.

### Allele frequencies of oxycodone metabolism enzymes in augmenter vs non-augmenter strains

Genetic variants (single nucleotide polymorphisms, SNPs) for the HRDP were sourced from a recent study ^25^. We identified 1,263 SNPs in the Cyp3a2, Cyp3a9, and Cyp2d1 genes (including exons, introns, and adjacent intergenic regions). In each sex, Fisher’s exact test was applied to compare the allele frequencies of all SNPs between Augmenters and Non-augmenters. The raw p-values were then adjusted using the false discovery rate.

### Estimate of heritability

The between-strain variance provides a measure of additive genetic variation (VA), while within-strain variance represents environment variability (VE). An estimate of narrow-sense heritability (i.e. the proportion of total phenotypic variation that is due to the additive effects of genes, *h*^2^) for nicotine or food reward was obtained using the formula: h2L=½*LVA/(½ *VA+VE) ^26,27^.

### Statistical analysis of behavioral data

The number of licks on active and inactive spouts was transformed into log scale to fit a normal distribution. The number of licks, reward, and intake during acquisition were analyzed by repeated measures ANOVA, where session and spouts were treated as within subject variables. PostHoc tests were conducted using the Tukey HSD procedure. Phenotypic correlations were determined using the Pearson test. Data were expressed as mean ± SEM. Statistical significance was assigned at p < 0.05. All analyses were conducted using R statistical language. Data are available on request from the authors.

## Acknowledgments

These studies were supported by NIDA U01DA053672 (B.M. Sharp, R.W. Williams and H. Chen) and by NIDA U01DA047638 (H. Chen and R.W. Williams). Breeders of the hybrid rat diversity panel were provided by Dr. Melinda R. Dwinell, Medical College of Wisconsin.

## Availability of Data

The data that support the findings of this study are available from the corresponding author upon reasonable request. Some data may not be available because of privacy or ethical restrictions.

## Supplementary Figures

**Figure S1.**
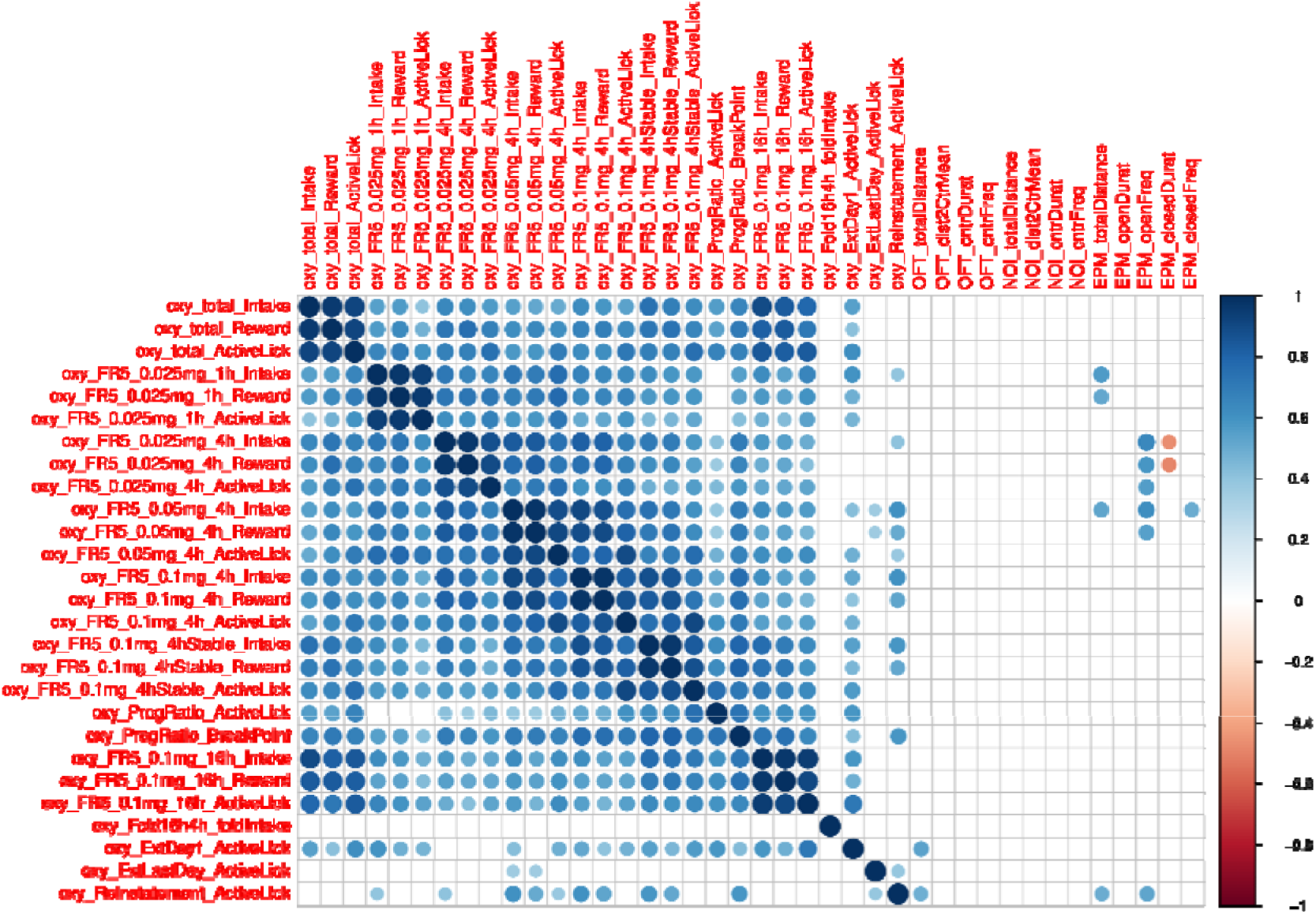
Correlation between oxycodone and behavioral traits in females.

**Figure S2.**
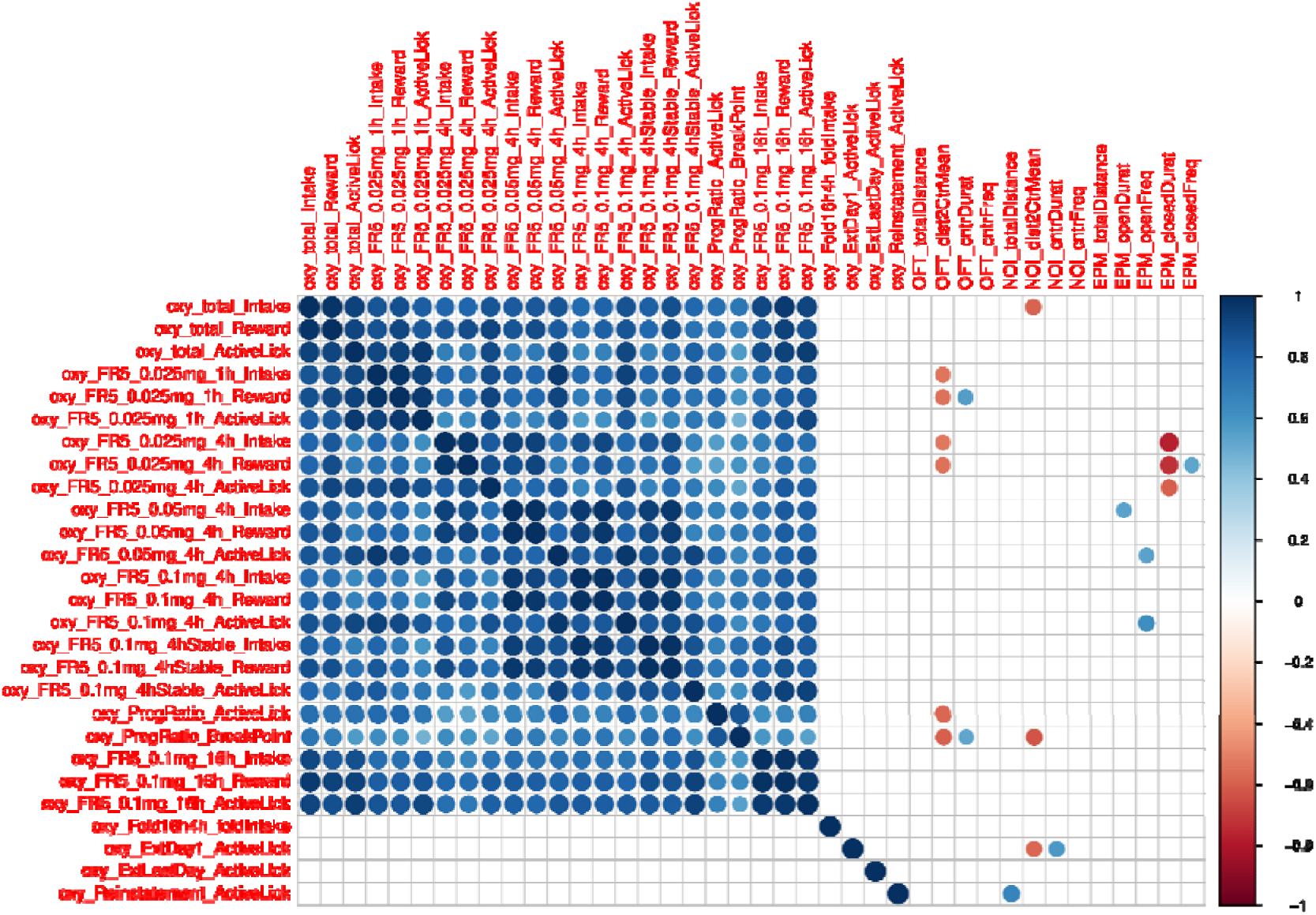
Correlation between oxycodone and behavioral traits in males.

